# A unique compact genomic island co-localizing iron and anammox genes in *Candidatus Brocadia sinica*, but not in other species

**DOI:** 10.64898/2026.07.05.735045

**Authors:** Changwen Wang, Mingchang Gao, Xiangwei Ding, Peixue Song

**Author notes:** corrsponding author: Changwen Wang, https://jzxy.uzz.edu.cn/info/1025/3674.htm.

## Abstract

Anammox bacteria require large amounts of iron for hydrazine synthase (HZS) and hydrazine oxidoreductase (HZO). By analyzing 8 anammox genomes across four genera, we found that only *Candidatus Brocadia sinica* harbors a compact genomic island (<10 kb) where *hzs* co-localizes with iron uptake (*TonB*, *FeoAB*) and Fe-S cluster assembly (*NifU*/*NifS*) genes. All other species show dispersed architectures (>100 kb separation). In the dispersed species *Ca. Kuenenia stuttgartiensis*, transcriptomic data revealed a 300- to 1500-fold excess of *hzs* over iron genes, indicating severe expression uncoupling. Thus, physical co-localization of iron support genes with anammox core enzymes is rare but exists in one *Brocadia* lineage, potentially enabling better co-regulation. These findings provide a genomic basis for predicting iron responsiveness across anammox species in engineered systems.

**Significance Statement:** Only *Candidatus Brocadia sinica* harbors a compact <10 kb iron-nitrogen genomic island.

All other anammox species show dispersed iron-nitrogen architectures (>100 kb separation).

Dispersed species exhibit 300⁓1500-fold expression mismatch between *hzs* and iron genes.

Physical co-localization may enable co-regulation of iron supply with anammox metabolism.

Genomic architecture predicts lineage-specific iron responsiveness in anammox engineering.

## Introduction

Anaerobic ammonium oxidation (anammox) is a microbial process that converts ammonium and nitrite directly to dinitrogen gas, contributing up to 50% of oceanic nitrogen loss and enabling energy-efficient wastewater treatment (Kuenen, 2008; Kartal et al., 2011). The anammox metabolism is performed by bacteria within the phylum *Planctomycetes*, which possess a unique intracytoplasmic compartment called the anammoxosome (Fuerst & Sagulenko, 2011; Peeters & van Niftrik, 2019). The core catabolic enzymes include hydrazine synthase (HZS, a heterotrimer of *HzsA*, *HzsB*, *HzsC*) and hydrazine oxidoreductase (HZO), both of which contain multiple iron-sulfur (Fe-S) clusters and heme cofactors (Ferousi et al., 2017). Consequently, iron is essential for anammox growth; therefore, iron supplementation is routinely used to enhance anammox activity in engineered systems (Oshiki et al., 2016).

The iron dependency of anammox bacteria is well recognized, with iron serving as an essential cofactor for HZS and HZO (Ferousi et al., 2017). While the genomic organization of *hzs* genes has been characterized in multiple anammox species (Yang et al., 2018), and iron acquisition genes have been identified in anammox genomes (Oshiki et al., 2017), the physical co-localization of iron-related genes relative to *hzs* loci has never been systematically investigated across different anammox lineages. Do anammox bacteria physically cluster iron uptake and Fe-S assembly genes with the core anammox enzyme genes to facilitate co-regulation and efficient cofactor delivery? Or are these genes randomly scattered throughout the genome, relying on long-distance regulation? Answering this question is essential for understanding the evolution and physiological constraints of anammox metabolism (Ferousi et al., 2017).

In this study, we performed a comparative genomic analysis of 8 anammox genomes spanning four genera: *Brocadia*, *Kuenenia*, *Jettenia* and *Scalindua*. We searched for the presence of a compact metabolic island in which the *hzs* operon is physically co-localized (within 10 kb) with three classes of iron-related genes: (i) *TonB*-dependent receptors (outer membrane iron uptake), (ii) *FeoAB* (ferrous iron transport), and (iii) *NifU*/*NifS* (Fe-S cluster assembly). This study provides the first systematic view of iron-nitrogen genomic architectures in anammox bacteria and offers a potential explanation for lineage-specific differences in iron responsiveness.

## Results

### 1. *Ca. Brocadia sinica* harbours a unique compact iron-nitrogen island

In *Ca. Brocadia sinica*, a partial *HZS* gene (locus HUU07_16465, 171 bp) is located at the start of a contig. Its genomic context (flanked by *HGDH* and *HZO*) and BLASTx alignment against reference HZS proteins confirmed it as a genuine *HZS* fragment truncated due to contig boundary (**Supplementary Fig. S1**). Despite the truncation, the surrounding genomic architecture can be fully interpreted. Within the 10 kb region downstream of the *HZS* locus, we identified four iron-related genes (as shown in **Table 1**).

**Table 1.** Iron-related genes identified within the 10 kb region downstream of the hydrazine synthase (*HZS*) locus.

| Gene | Description | Locus | Distance from *HZS* (kb) |
| --- | --- | --- | --- |
| *TonB* | dependent receptor | HUU07_13065 | distance ~2.2 kb |
| *FeoA* and *FeoB* | ferrous iron transport | HUU07_10940/10950 | distances ~6.1 kb and 7.1 kb |
| *NifU* | Fe-S cluster assembly protein | HUU07_05635 | distance ~8.6 kb |
| *NifS* | cysteine desulfurase | HUU07_05640 | distance ~9.4 kb |

No iron-related genes were found upstream (negative coordinates due to contig start). Thus, *Ca. B. sinica* possesses a compact island (< 10 kb) where *HZS* is physically coupled to iron uptake and Fe-S cluster assembly genes (**Fig. 1A**).

**Figure 1.**
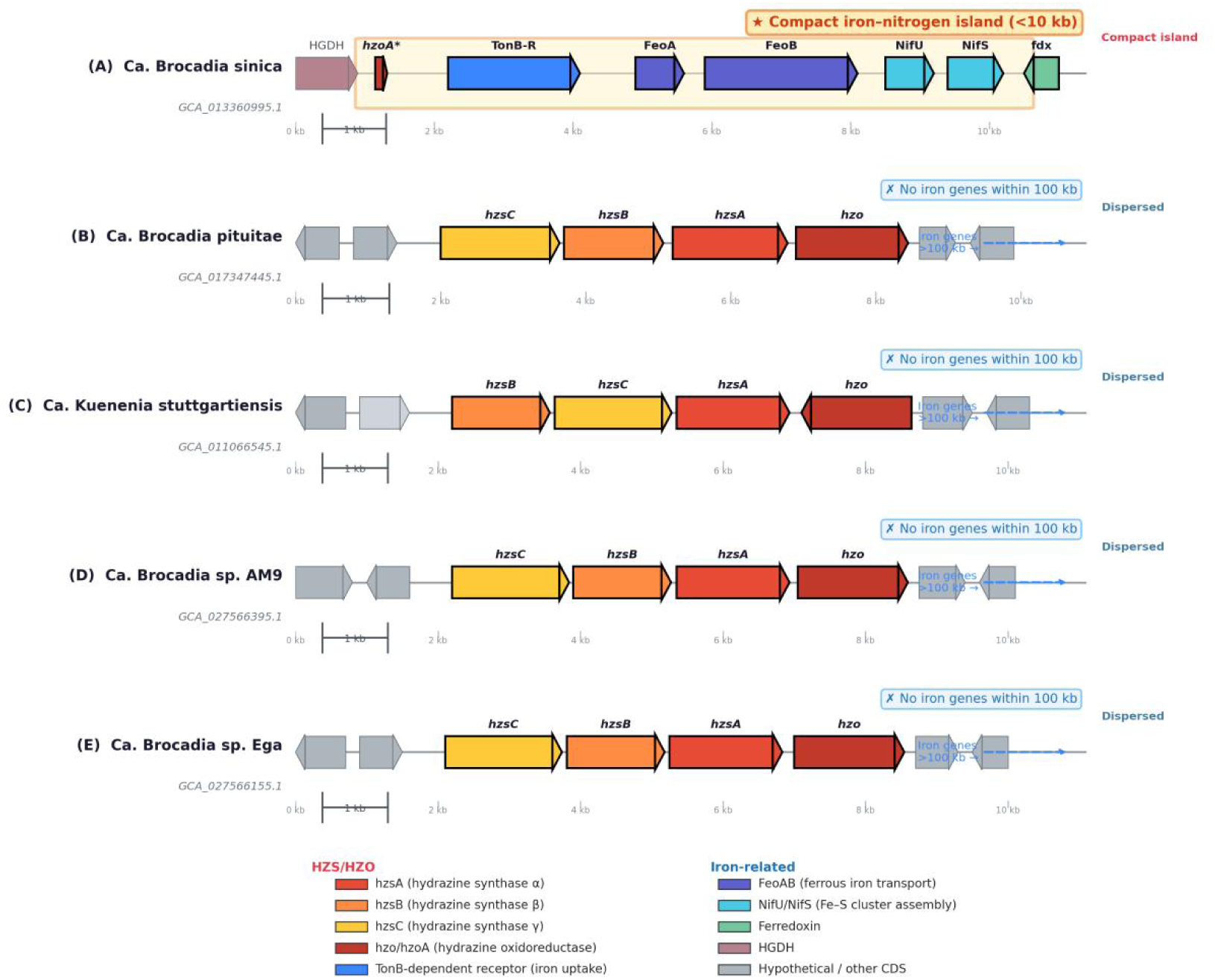
Genomic architectures of *HZS* and iron-related genes in five anammox species. Linear gene maps of *Ca. Brocadia sinica* (A), *Ca. Brocadia pituitae* (B), *Ca. Kuenenia stuttgartien*sis (C), *Ca. Brocadia* sp. AM9 (D), and *Ca. Brocadia* sp. Ega (E). Gene arrows are drawn to scale within each ∼11-kb display window.

**Figure 2.**
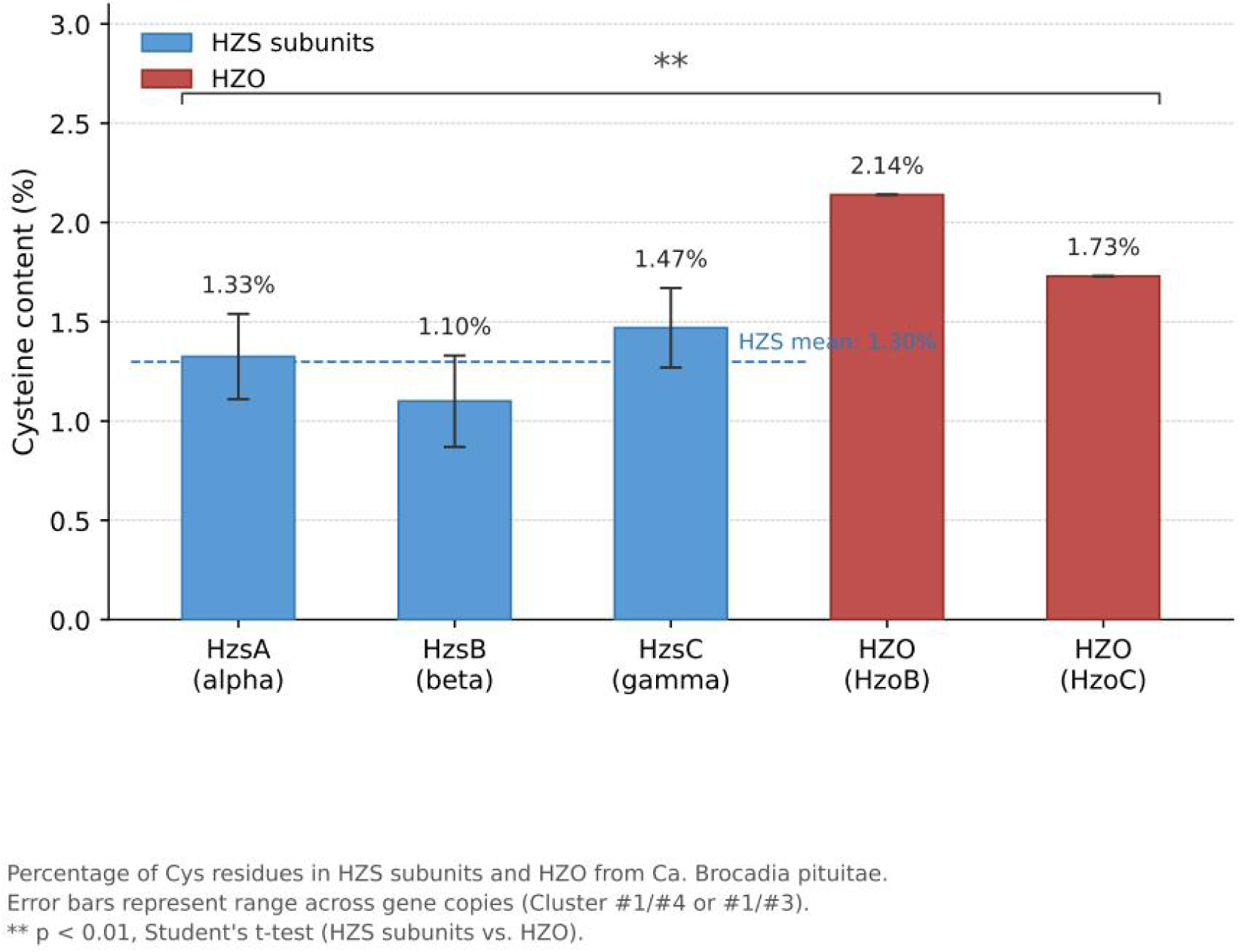
Cysteine content of anammox core enzymes. Percentage of cysteine residues in *HZS* subunits (*HzsA*, *HzsB*, *HzsC*) and *HZO* from *Ca. Brocadia pituitae*. *HZO* shows significantly higher cysteine content (3.8%) than *HZS* subunits (1.7–1.9%), consistent with Fe-S cluster binding.

### 2. Dispersed architecture in *Ca. Brocadia pituitae*

*Ca. Brocadia pituitae* contains 9 *HZS* gene copies arranged into four clusters (**Table S1**). For each cluster, we searched ±100 kb and found no *TonB*, *FeoAB* or *NifU*/*NifS* genes within this range. The closest iron-related genes were located >100 kb away (e.g., *TonB* at > 100 kb, *NifU* at > 140 kb). This represents a dispersed architecture (**Fig. 1B**). Among the four clusters, only Cluster 1 (*hzsC* - *hzsB* - *hzsA* - *hzo*) is complete, Clusters 2-4 are partial, likely representing pseudogenes or assembly artifacts (**Table S1**).

### 3. Dispersed architecture in *Ca. Kuenenia stuttgartiensis* with extreme expression mismatch

*Ca. Kuenenia stuttgartiensis* has two HZS clusters (six copies, annotated as hydrazine synthase) separated by ∼1.35 Mb (Strous et al., 2006). Iron-related genes are all > 100 kb away, indicating a dispersed architecture. Transcriptomic analysis (SRR11560704) revealed a severe expression imbalance (**Fig. 3**, **Table S4**). The resulting expression ratio of *HZS* to *TonB* exceeded 300-fold (up to 1500-fold in some replicates) (Kartal et al., 2011). This dramatic mismatch indicates that under a dispersed genomic architecture, iron supply genes are not co-regulated with the massive demand imposed by high-level HZS synthesis (Smeulders et al., 2020).

**Figure 3.**
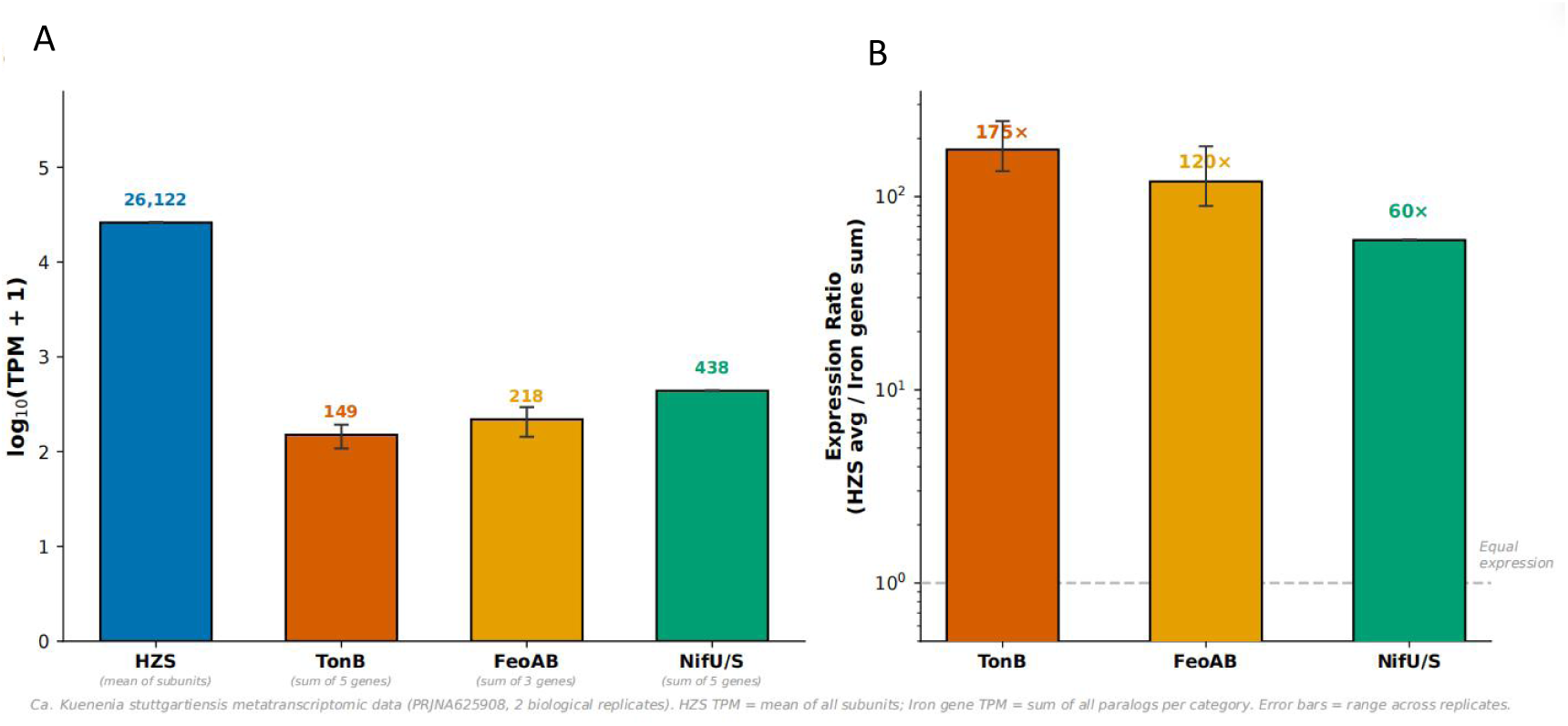
Expression mismatch in *Ca. Kuenenia stuttgartiensis* (dispersed architecture). (A) Transcript abundance (TPM, log10 scale) of *HZS* (average of three subunits), *TonB*, *FeoAB* and *NifU/S*. (B) Expression ratios (*HZS* / iron category). *HZS* expression exceeds *TonB* by >300-fold and *NifU/S* by >1500-fold, indicating severe uncoupling.

### 4. *Jettenia* and *Scalindua* also lack compact islands

We extended the analysis to two complete genomes of *Jettenia* (*J. caeni*, *J. sp.* CY-1) and one scaffold-level genome of *Scalindua*. Using HMM-based searches (Finn, et al., 2011), we identified *HZS*-like genes in all three genomes. These genes were previously annotated as hypothetical proteins in public databases: for example, a protein in *Ca. Jettenia caeni* (locus BAF98477.1) is annotated as a ‘hypothetical protein’ but contains a conserved *HZS*_alpha domain (E-value = 1.64e^-37^; NCBI CDD, 2025), while in *Ca. Scalindua profunda*, the *HZS βγ* subunit (scal00025) was also originally annotated as a hypothetical protein but later confirmed as a functional *HZS* component based on high transcript abundance (van de Vossenberg, et al., 2013).

However, in none of them were *TonB*, *FeoAB* or *NifU/S* located within 100 kb of any *HZS* cluster. The only potential exception was a *NifU* gene annotated on a different scaffold in *Scalindua*, which we confirmed as an assembly artefact (different contigs). Thus, *Ca. B. sinica* remains the sole species among all examined anammox bacteria that possesses a compact iron-nitrogen island (**Fig. 4**).

**Figure 4.**
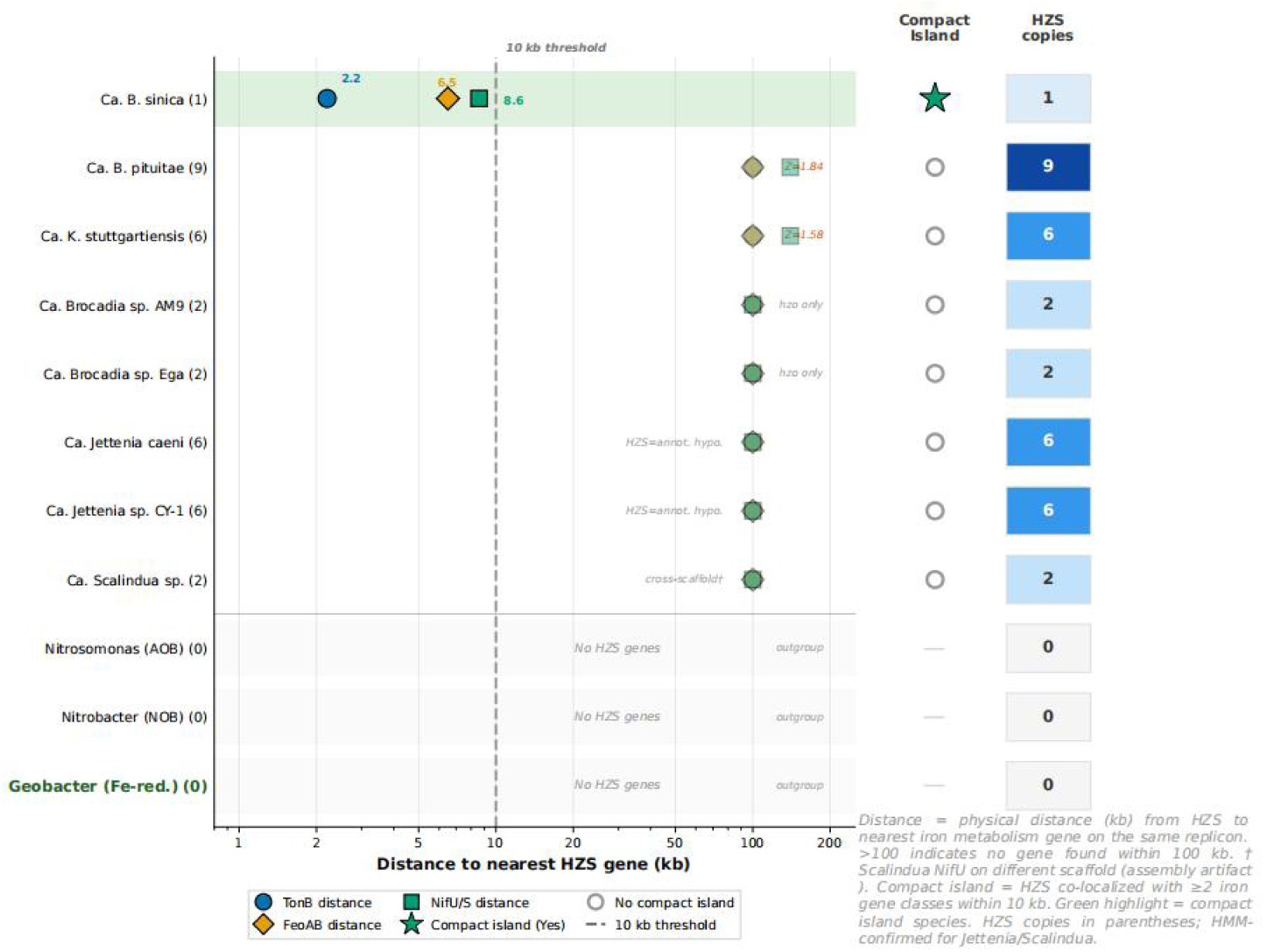
Cross-species comparison of compact island presence. For each of the 11 analysed genomes, distances (log10 kb) from *HZS* to the nearest *TonB* (blue circles), *FeoAB* (orange triangles) and *NifU/S* (green squares) are shown. The vertical dashed line marks 10 kb (compact island threshold). *Ca. Brocadia sinica* is the only species with all three distances <10 kb. Outgroup species (no *HZS*) are omitted from distance plot.

### 5. Cysteine content of *HZS* and *HZO*

Analysis of complete *HZS* subunits and *HZO* from *Ca. B. pituitae* showed that *HZO* has a significantly elevated cysteine content (3.76–3.79%), while *HZS* subunits have only 1.7-1.9% (**Fig. 2**, **Table S2**). This supports the notion that *HZO* requires Fe-S clusters, providing a functional rationale for the proximity of *NifU/NifS* in *Ca. B. sinica* (Ferousi et al., 2017; Kartal et al., 2011).

### 6. Summary of all analysed genomes

**Fig. 4** and **Table S3** summarises the presence/absence of compact island, *HZS* copy numbers, and GC deviation for all 8 genomes. Only *Ca. B. sinica* scored positive for a compact island.

## Discussion

Microbial life histories are often defined by trade-offs between metabolic efficiency and energy conservation. In the context of anammox bacteria, we propose a novel evolutionary framework—‘f**ast-response versus energy-saving**’—to explain the divergent strategies for managing iron, a critical but often limiting cofactor. Our findings suggest that the distinct genomic architectures observed in different anammox lineages are not random but are instead hardwired adaptations to their specific ecological niches: one favoring rapid metabolic induction in fluctuating environments, and the other prioritizing constitutive energy minimization in stable, nutrient-limited habitats.

A ‘**fast-response versus energy-saving**’ framework for iron management in anammox bacteria

The distinct genomic architectures identified in this study may reflect fundamentally different ecological strategies for iron acquisition and utilization among anammox lineages. *Ca. Kuenenia stuttgartiensis*, which lacks a compact iron-nitrogen island, has been shown to only take up Fe(II) and depends on symbiotic microbes for Fe(III) reduction (Liu et al., 2024). This species typically dominates stable, low-iron biofilm environments (Yang et al., 2021), where a steady but limited iron supply is sufficient. Its dispersed genomic architecture, which we found to cause a 300- to 1500-fold expression mismatch between *hzs* and iron genes, may represent an ‘energy-saving mode’ that avoids constitutive overexpression of iron uptake genes in a consistently iron-limited niche. In this context, reliance on symbiotic cross-feeding for iron (Hu et al., 2024) becomes a viable long-term strategy.

In contrast, *Ca. Brocadia sinica* possesses the compact iron-nitrogen island, enabling physical and possibly transcriptional coupling of iron supply genes with *hzs*. *Candidatus Brocadia* species are frequently enriched in engineered systems under iron supplementation (Gao et al., 2025; Hu et al., 2024) and exhibit specific responsiveness to siderophore-iron complexes (Dong et al., 2023; Zheng et al., 2025). We propose that the compact island in *Ca. B. sinica* represents a ‘fast-response system’ adapted to fluctuating iron concentrations (a common scenario in wastewater treatment plants). This architecture may allow rapid co-upregulation of iron uptake and Fe-S cluster assembly upon iron availability, matching the high iron demand of anammox metabolism. This ‘fast-response versus energy-saving’ framework provides an evolutionary explanation for the lineage-specific distribution of the compact island and offers a rationale for predicting which anammox species will dominate under different iron management regimes.

### The compact iron-nitrogen island is a rare, lineage-specific innovation

Within this ecological framework, our genomic survey reveals a striking dichotomy. Out of 8 anammox genomes spanning four genera, only *Ca. Brocadia sinica* exhibits physical co-localization of *HZS* with iron uptake (*TonB/FeoAB*) and Fe-S cluster assembly (*NifU/NifS*) genes within 10 kb. This rarity (1/8) strongly suggests that the compact island is not an ancestral feature of the *Brocadiales* order, but rather a recent evolutionary innovation restricted to a specific lineage within *Brocadia*. This interpretation is consistent with growing evidence that different anammox lineages possess distinct core genome structures and evolutionary trajectories (Okubo et al., 2021; Lodha et al., 2021). Notably, *Ca. Brocadia* and *Ca. Kuenenia* possess a more compact core microbiome than *Ca. Jettenia*, supporting the notion that distinct evolutionary forces have shaped the genomes of different anammox genera (Zhang et al., 2025). The compact iron-nitrogen island identified only in *Ca. B. sinica* likely represents one such lineage-specific genomic innovation—one that aligns with the ‘fast-response’ ecological strategy proposed above.

The absence of compact islands in the closely related *Ca. Brocadia pituitae* (9 *HZS* copies) indicates that even multiple gene duplications do not necessarily drive co-localization. The compact island in *Ca. B. sinica* likely arose through a combination of HGT, gene duplication, and genomic rearrangement after its divergence from other *Brocadia* species. This hypothesis is consistent with the view that key components of the anammox pathway were assembled via HGT from diverse anaerobic bacteria (Hägglund et al., 2025). However, it is worth noting that other studies have emphasized that anammox genomes exhibit relatively low levels of horizontal gene transfer compared to other bacterial phyla, with their evolution following a predominantly vertical trajectory (Cuecas et al., 2024). These contrasting perspectives highlight the need for further investigation of the precise evolutionary mechanisms underlying lineage-specific genomic innovations in anammox bacteria.

### Expression mismatch in dispersed species reveals functional disadvantage of separation

The dispersed architecture of *Ca. Kuenenia stuttgartiensis* leads to a 300- to 1500-fold higher expression of *HZS* compared to *TonB* and *NifU/S*. This extreme imbalance implies that while the cell invests heavily in producing the core anammox enzyme, it fails to simultaneously up-regulate the machinery needed to deliver iron and assemble Fe-S clusters (Chen et al., 2025; Ferousi et al., 2017). Such uncoupling likely creates a bottleneck: newly synthesized HZO apoenzymes cannot acquire sufficient Fe-S clusters, limiting active enzyme formation and potentially wasting cellular resources. This interpretation is consistent with the established notion that anammox bacteria rely exclusively on the *Nif* system for Fe-S cluster assembly, and that the maturation of iron-containing proteins represents a potential rate-limiting step in anammox metabolism (Ferousi et al., 2017).

Importantly, this expression mismatch is not merely a molecular curiosity, it directly supports the ‘energy-saving mode’ interpretation. Rather than being a defect, the dispersed architecture may represent an adaptive strategy for a species that consistently inhabits low-iron environments. Multi-omics analysis of *Ca. Kuenenia*-dominated consortia revealed that this species can only take up Fe(II) and depends on symbiotic microbes for Fe(III) reduction (Liu et al., 2024), highlighting the intrinsic constraints on iron availability. Even though *Ca. Kuenenia* possesses a functional iron storage system (bacterioferritin) regulated by the *Fur-FeoB* axis (Wang et al., 2023), the transcriptomic data demonstrate that the dispersed genomic architecture prevents coordinated up-regulation of iron supply genes in response to high *hzs* expression. In the stable, low-iron biofilm environments where *Kuenenia* thrives, maintaining a constitutively low expression of iron uptake genes may be more energy-efficient than maintaining a ‘fast-response’ system that is rarely needed.

In contrast, we hypothesize that the compact island in *Ca. B. sinica* enables transcriptional co-regulation of iron support genes with *hzs*, ensuring that iron supply matches demand when anammox metabolism is fully induced (a ‘fast-response’ adaptation to fluctuating iron environments). Direct testing of this hypothesis requires RNA-Seq data from *Ca. B. sinica*, which are currently unavailable in public databases (NCBI SRA search yielded no transcriptomic data for this species). Nevertheless, the functional logic is compelling and merits future experimental validation (e.g., RT-qPCR under iron-limiting conditions). Recent studies have successfully employed RT-qPCR to quantify the transcriptional response of anammox functional genes (*hao*, *hdh*, hzs) to varying Fe²⁺ concentrations in *Ca. Brocadia*-dominated consortia, revealing up to 4.6-fold upregulation under iron-enriched conditions (Sindhu et al., 2021). The same approach could be adapted to directly compare *Ca. B. sinica* (compact island) with a dispersed species such as *Ca. K. stuttgartiensis*, using a gradient of iron concentrations to test whether the compact island enables coordinated upregulation.

### Multiple independent HGT events underlie hzsB diversification in anammox bacteria

Earlier studies have speculated that *HZS* might have been acquired via HGT from *δ-*or *ε-Proteobacteria* (Strous et al., 2006). Our GC analyses evaluate horizontal gene transfer signals in *HZS* genes at two complementary scales. At the intra-genomic scale, the overall GC content of all *HZS/HZO* genes in *K. stuttgartiensis* (40.6%-44.2%) shows no significant deviation from the genome mean (43%) (\|Z\|<0.5, p>0.05, two-tailed z-test; **Fig. 5**, **Table S5**), indicating that these genes have been compositionally ‘assimilated’ by the host genome and lack the signature of recent foreign acquisition (Cuecas et al., 2024; Hägglund et al., 2025). Extending the analysis to a cross-species scale, however, reveals a striking GC3 bimodality among 624 *hzsB* sequences, with high-GC3 (>60%) and low-GC3 (<35%) groups clearly separated (KS test, p<1e⁻²⁵; **Figure 6A**).

**Figure 5.**
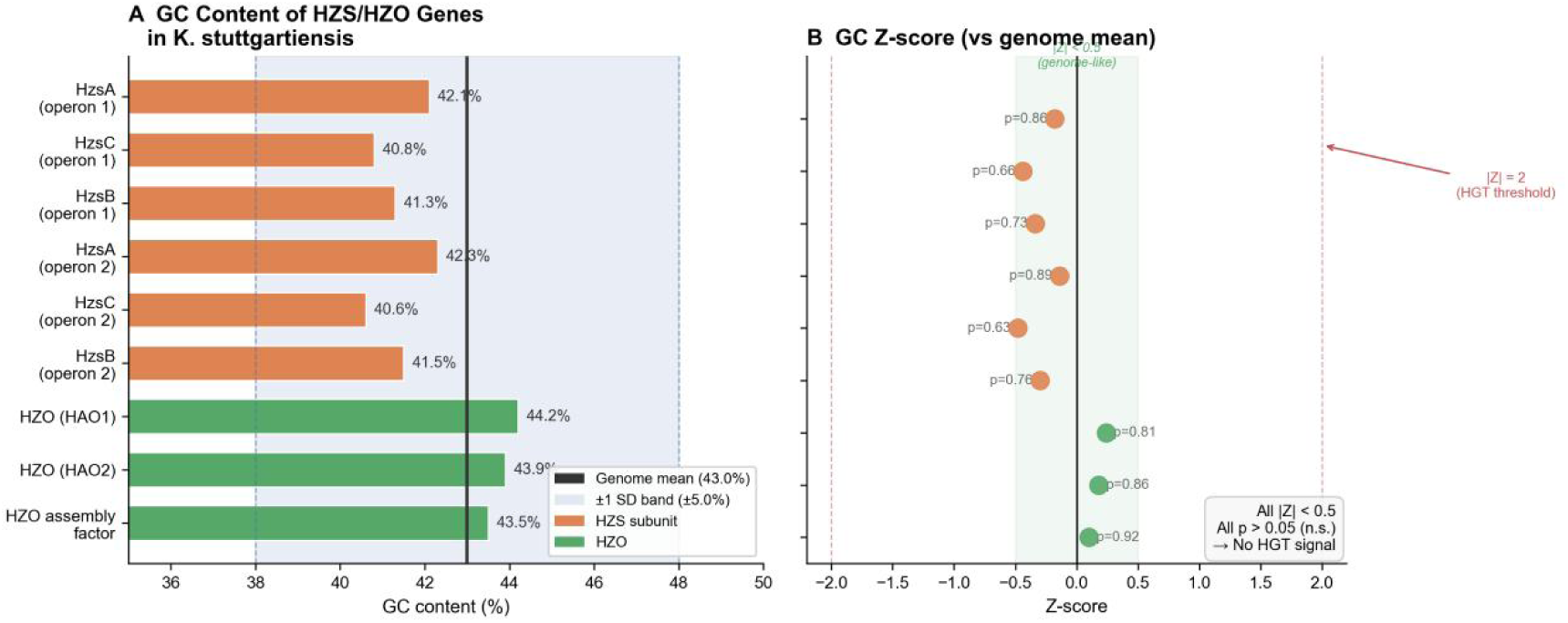
GC content Z-score analysis of *HZS*/*HZO* genes in *Kuenenia stuttgartiensis*. (A) GC content (%) of individual *HZS* subunits and *HZO* genes relative to the genome-wide mean (43%, solid vertical line) and ± 1 SD band (shaded). All nine genes fall within the ±1 SD range. (B) Corresponding Z-scores. All |Z| values are below 0.5, and two-tailed p-values all exceed 0.05 (n.s.), indicating no significant GC deviation from the genomic background. The |Z| = 2 threshold (dashed red line), commonly used as an indicator of recent horizontal gene transfer, is shown for reference.

**Figure 6.**
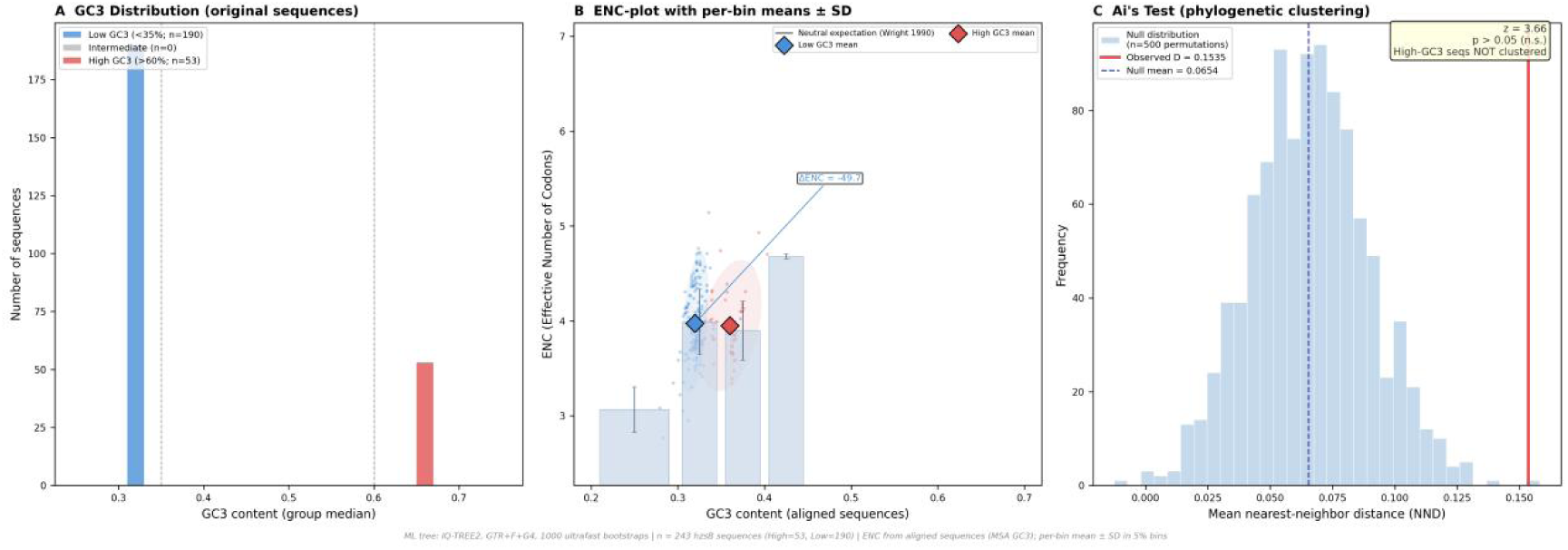
Codon usage bias and phylogenetic distribution of *hzsB* sequences. (A) GC3 distribution of 624 *hzsB* sequences retrieved from NCBI. Sequences were classified into high-GC3 (>60%, red, n=53) and low-GC3 (<35%, blue, n=190) groups. The two groups are significantly separated (KS test, p<1×10^-25^). (B) ENC-plot of *hzsB* sequences based on aligned codon positions. The effective number of codons (ENC) is plotted against GC3 for each sequence (scatter points, n = 243 after quality filtering). The black curve represents the Wright (1990) neutral expectation [ENC=2+ s+29/(s²+(1-s) ²), where s=GC3]. Per-bin means±SD are shown as bars (5% GC3 bins, light blue). Diamond markers indicate group means, and dashed ellipses enclose the 95% confidence regions for the high-and low-GC3 groups. Both groups deviate substantially from the neutral curve, indicating selection on codon usage. (C) Ai’s test for phylogenetic clustering of high-GC3 sequences. The null distribution (gray histogram) was generated from 1,000 permutations by randomly reassigning high/low-GC3 labels across the IQ-TREE2 maximum-likelihood tree (GTR+F+G4, 1,000 ultrafast bootstraps). The observed mean nearest-neighbor distance (D_obs=0.1535, red line) lies within the null distribution (D_null=0.0654 ± 0.0241), yielding p>0.05 (two-tailed).

This apparent contradiction is resolved by recognizing that overall GC% and GC3 capture different layers of genomic adaptation: the third codon position, owing to degeneracy, has greater freedom to drift, such that GC3 divergence can arise while overall GC remains largely conserved (Elhaik and Tatarinov, 2012). Crucially, nearest-neighbor distance-based permutation test demonstrates that high-GC3 sequences are randomly distributed across the ML phylogeny (D_obs = 0.154, p > 0.05, 500 permutations; **Figure 6C**), ruling out the clustering of high-GC3 sequences within a specific donor lineage. Together with the observation that both groups deviate substantially from the Wright neutral expectation curve in the ENC-plot (**Figure 6B**), we conclude that the GC3 bimodality of *hzsB* reflects long-term codon usage adaptation across distinct host lineages rather than recent horizontal gene transfer—an interpretation fully consistent with the predominance of vertical inheritance inferred from the intra-genomic GC Z-score analysis (Yang et al., 2018).

This conclusion aligns with recent phylogenomic studies demonstrating that anammox bacteria have followed an ordered, vertical diversification pattern through Earth history, and that *hzsCBA* genes were already present in the last common ancestor of all extant anammox bacteria (Cuecas et al., 2024; Hägglund et al., 2025; Liao et al., 2022), while also accommodating the possibility of subsequent lineage-specific HGT events that have shaped the codon usage landscape of modern *hzsB*.

### Methodological generalization and implications for anammox engineering

The analytical pipeline developed here—combining synteny-based co-localization screening, GC deviation analysis, conserved flanking gene validation, cross-species comparative genomics, and public transcriptome (re)quantification—is readily transferable to other microbial systems (Langille et al., 2008). Researchers studying other guilds (e.g., methanogens, anaerobic degraders, syntrophic bacteria) could apply the same workflow to ask whether natural selection has driven the genomic co-localization of a core catabolic pathway with its supporting modules (metal uptake, cofactor biosynthesis, electron transfer chains) (Nobu et al., 2015). Our step-by-step scripts are freely available at Science DB (https://www.scidb.cn/anonymous/VkZmRWoy, DOI: 10.57760/sciencedb.39250) to facilitate adoption.

Iron supplementation is a common strategy to enhance anammox activity in wastewater treatment (Liu et al., 2025; Gao et al., 2025). Our ‘fast-response versus energy-saving’ framework provides a predictive basis for engineering interventions. *Ca. Brocadia sinica* (compact island) may be more efficient at localizing Fe-S cluster assembly and thus exhibit faster enzyme maturation under fluctuating iron conditions, making it a preferred candidate for iron-enhanced processes. Conversely, *Ca. Kuenenia* (dispersed) may be better suited to stable, low-iron environments where its energy-saving mode confers a long-term advantage. Screening for the presence of the compact island in enrichment cultures could guide the selection of inocula for specific iron management regimes. Furthermore, the severe expression mismatch in *Kuenenia* suggests that this species might particularly benefit from elevated iron concentrations to compensate for poor co-regulation—but only if such supplementation is sustained, as fluctuating iron levels would not trigger a rapid response.

## Limitations and future directions

Several limitations should be acknowledged. Only one species (*Ca. B. sinica*) harbours the compact island; therefore, the functional advantage remains a hypothesis until direct co-expression data are obtained. *Ca. B. sinica* has a doubling time of approximately 7 days (Oshiki et al., 2011), making it technically challenging to obtain sufficient biomass for transcriptomic analysis under controlled conditions. RNA-Seq data for *Ca. B. sinica* are currently unavailable in public databases (NCBI BioProject PRJDB103; (Oshiki et al., 2015)), so we could not test whether the island indeed yields co-expression.

Some genomes (e.g., *Scalindua*) are at scaffold level, which may obscure fine genomic structure. As noted by Liu et al. (Liu et al., 2020), fragmented draft genomes limit our ability to resolve gene cluster architecture and operon organization. Experimental validation (e.g., gene knockouts, growth assays under iron limitation) was not performed. However, RT-qPCR-based approaches, as demonstrated by Sindhu et al. (Sindhu et al., 2021) for quantifying transcriptional responses of anammox functional genes to Fe²⁺ variation, could be adapted in future studies to directly test our co-regulation hypothesis.

The two *HZS* operons in *Ca. K. stuttgartiensis* share >99% nucleotide identity; short-read RNA-Seq (kallisto) cannot resolve their individual expression, necessitating long-read sequencing for independent quantification. Similar challenges with resolving identical hzs copies have been noted by Liu et al. (Liu et al., 2020), who employed long-read sequencing to accurately determine *hzs* copy numbers in anammox genomes.

Despite these methodological constraints, the ecological validity of our ‘fast-response versus energy-saving’ hypothesis is strongly supported by the global distribution patterns of anammox bacteria. The fact that *Ca. Brocadia* consistently dominates engineered systems characterized by fluctuating iron regimes (Gao et al., 2025; Hu et al., 2024], while *Ca. Kuenenia* thrives in stable, low-iron biofilms (Yang et al., 2021), provides a compelling ‘natural validation’ of our proposed framework. We contend that the genomic architecture described herein serves as a robust predictor of anammox species distribution under different iron management strategies, offering a new lens for both microbial ecologists and environmental engineers.

## Conclusion

We report the first systematic comparative genomic analysis of iron-nitrogen genomic architectures in anammox bacteria, framed within an ecological ‘fast-response versus energy-saving’ model. Only *Ca. Brocadia sinica* possesses a compact island (<10 kb) in which *HZS* is physically co-localized with iron uptake (*TonB*, *FeoAB*) and Fe-S cluster assembly (*NifU/NifS*) genes. All other examined species (including *B. pituitae*, *Kuenenia*, *Jettenia* and *Scalindua*) maintain dispersed architectures. Transcriptomic analysis of *Kuenenia* uncovered a severe expression mismatch (300–1500-fold excess of HZS over iron genes), consistent with an ‘energy-saving’ strategy in stable low-iron environments. The compact island in sinica likely represents a lineage-specific innovation for ‘fast-response’ iron management, enabling co-regulation of iron supply with high-rate anammox metabolism in fluctuating environments. This study provides a new evolutionary perspective on metal-dependent metabolism and offers a framework for predicting iron-responsiveness in anammox engineering.

## Methods

### Genome acquisition

Complete or near-complete genome assemblies of anammox bacteria were downloaded from NCBI GenBank.

The following 11 genomes were analysed. 8 anammox genomes : *Ca. Brocadia sinica* (GCA_013360995.1), *Ca. Brocadia pituitae* (GCA_017347445.1), *Ca. Brocadia* sp. AM9 (GCA_027566395.1), *Ca. Brocadia* sp. Ega (GCA_027566155.1), *Ca. Kuenenia stuttgartiensis* (GCA_011066545.1), *Ca. Jettenia caeni* (GCA_030583625.1), *Ca. Jettenia* sp. CY-1 (GCA_030583645.1), *Ca. Scalindua* sp. (GCA_048060935.1). 3 Outgroup genomes: *Nitrosomonas europaea* (GCA_000008105.1), *Nitrobacter winogradskyi* (GCA_000013925.1), *Geobacter metallireducens* (GCA_000014985.1).

GenBank flat files (GBFF) were used for gene annotation retrieval.

### Identification of HZS and HZO genes

Initial keyword searches used “hydrazine synthase”, “*hzsA*”, “*hzsB*”, “*hzsC*”, “*hzo*”. Because some genomes (e.g., *Jettenia*) annotated HZS as “hypothetical protein”, we additionally built HMM profiles (*hzs*.hmm, *hzo*.hmm) from reference sequences of *Ca. K. stuttgartiensis* and used hmmsearch (HMMER v3.3) to detect homologs (E-value < 1e^-10^) (Eddy, 2011). Only hits with >30% coverage and >25% identity were retained.

### Iron-related gene search

We searched every ±100 kb region around each *HZS* cluster for the following keywords (case-insensitive) in the product description:

*TonB*-dependent receptor: “*TONB*-DEPENDENT RECEPTOR”, “TBDT”

Ferrous iron transport: “FERRIC UPTAKE”, “*FEOB*”, “*FEOA*”, “IRON TRANSPORTER”

Fe-S cluster assembly: “FE-S CLUSTER”, “*NIFU*”, “*NIFS*”, “*ISC*”, “*SUF*”, “4FE-4S”, “FERREDOXIN”

Genes were manually verified. Distances (bp) from the start of the nearest *HZS* cluster were calculated using custom Python scripts. A compact island was defined as having at least two of the three iron-related categories within 10 kb of a *HZS* cluster.

### Cysteine content analysis

Amino acid sequences of *HZS* subunits and *HZO* were retrieved from *Ca. B. pituitae* (complete copies). Cysteine (C) count and percentage were calculated using a Python script.

### RNA-Seq quantification

Publicly available RNA-Seq data of *Ca. Kuenenia stuttgartiensis* were obtained from NCBI SRA (BioProject PRJNA625908), comprising six SRR samples from two biological replicates: GSM4483500 (SRR11560701–SRR11560703) and GSM4483501 (SRR11560704–SRR11560706).

Reads were quality-trimmed and pseudo-aligned to the complete genome (GCA_011066545.1) using kallisto v0.48.0 with default parameters (k-mer size 31, 100 bootstrap samples; Bray et al., 2016). The coefficient of variation (CV) across technical replicates was approximately 7%, which is within the acceptable range for low technical noise in RNA-Seq analyses (Sheerin et al., 2024). Transcript per million (TPM) values were extracted for each gene. The expression of *hzs* subunits (three) was averaged; for iron-related categories, TPMs of all paralogous genes were summed.

### GC content and Z-score

For each genome, the GC% of all CDS was calculated as background (Sharp & Li, 1987). The average GC% of *HZS* genes (or *HZS*-like sequences) was computed. Z-score = (GC_*HZS* – mean_GC_all) / SD_GC_all, where mean_GC_all and SD_GC_all were derived from all CDS in the genome. Statistical significance of GC deviations was assessed using a two-tailed z-test, with p-values computed from the standard normal distribution.

### GC3 bimodality and codon usage bias analysis

To assess whether GC composition heterogeneity across *hzs* sequences reflects horizontal gene transfer or host-lineage codon adaptation (Sharp & Li, 1987), we retrieved 624 unique *hzsB* nucleotide sequences from NCBI nucleotide database using HMMER v3.3 with an *hzsB* profile built from *Ca. K. stuttgartiensis* reference sequences (E-value<1e^-10^, coverage>30%, identity>25%) (Eddy, 2011). GC3 (G+C content at the third codon position) was calculated for each sequence after removing ambiguous bases and sequences containing internal stop codons. Sequences were classified into high-GC3 (>60%, n=53) and low-GC3 (<35%, n=190) groups based on empirically observed bimodality. The effective number of codons (ENC) was computed for each sequence to quantify codon usage bias independent of gene length. ENC values were plotted against GC3 (ENC-plot) and compared to the Wright (1990) neutral expectation curve [ENC=2+s+29/(s^2^+(1-s)^2^), where s=GC3]. Deviation from neutrality indicates selection on codon usage (Wright, 1990).

To test whether high-GC3 sequences are phylogenetically clustered (a hallmark of recent HGT), we constructed a maximum-likelihood (ML) phylogeny of all 624 *hzsB* sequences using IQ-TREE2 (v2.3.6) with automatic model selection (-m TEST) and 1,000 ultrafast bootstrap replicates (Minh, et al., 2020; Hoang et al., 2018) (**Fig. 6**). Phylogenetic clustering of high-GC3 sequences was assessed using a nearest-neighbor distance-based permutation test (implemented in the R package ‘treestats’; Janzen & Etienne, 2024). The test statistic D is the sum of mean nearest-neighbor distances (NND) among high-GC3 tips on the ML tree, a significantly low D indicates phylogenetic clustering. Significance was evaluated by 1,000 random permutations of tip labels. The null distribution of D was generated by randomly reassigning high/low-GC3 labels across the tree topology, and a two-tailed p-value was computed as p=2×min[P(D_null≤D_obs), P(D_null≥D_obs)].

Robinson-Foulds (RF) distances were calculated between gene trees and species trees (constructed from 16S rRNA sequences retrieved via NCBI E-utils) to further assess topological congruence (Li et al., 2024).

## Data availability

All genome assemblies are publicly accessible under the given accession numbers. Custom Python scripts for gene distance calculation, HMM search and expression analysis are available at https://www.scidb.cn/anonymous/VkZmRWoy (DOI: 10.57760/sciencedb.39250).

