## Supplementary figures for "A unique compact genomic island co-localizing iron and anammox genes in *Candidatus Brocadia sinica*, but not in other species"

**Supplementary Figure**

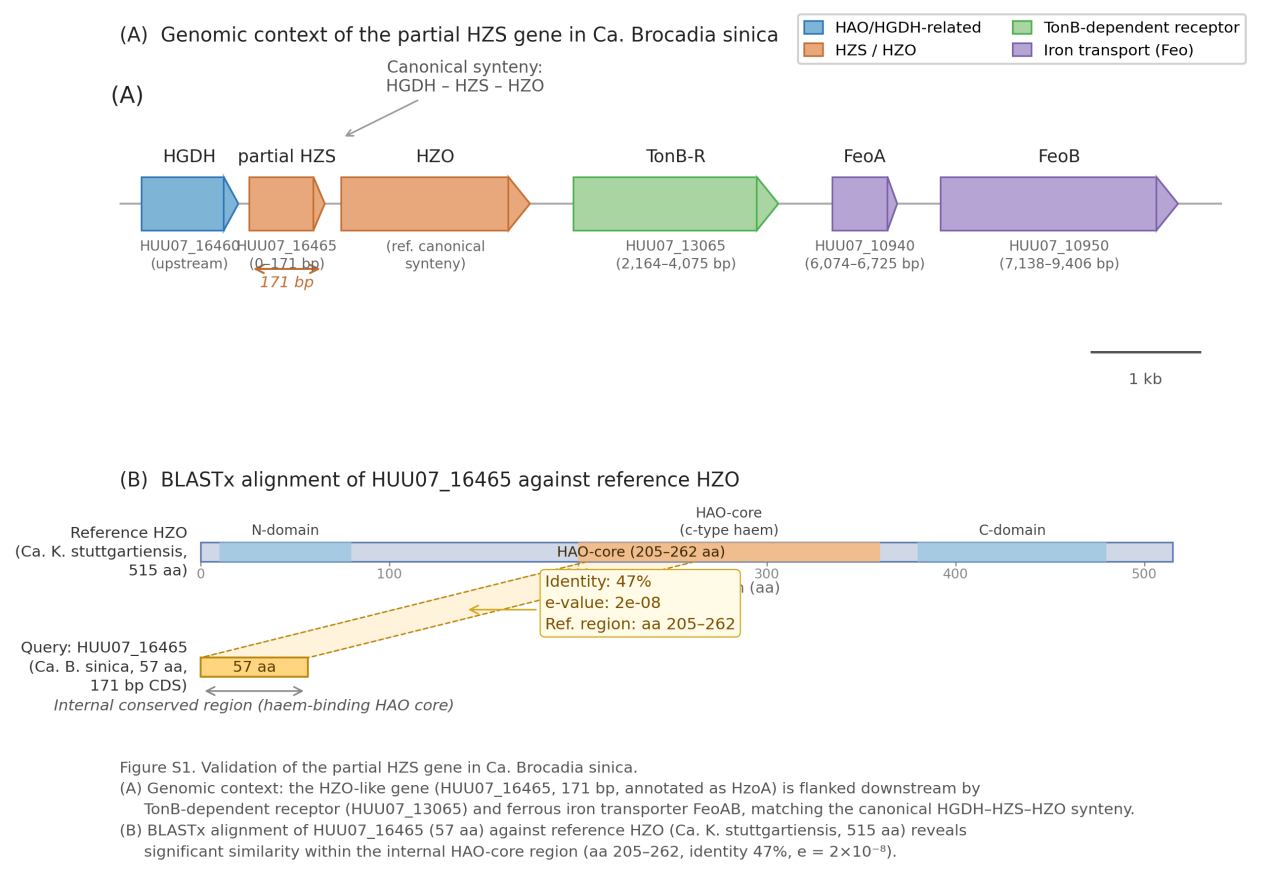

**Supplementary Figure S1.** **Validation of the partial *HZS* gene in Ca. Brocadia sinica. (A) Genomic context: *HGDH*-partial *HZS*-*HZO*, matching the canonical synteny. (B) BLASTx alignment against reference *HZO*, showing internal conserved region.**
