## Supplementary table for "A unique compact genomic island co-localizing iron and anammox genes in *Candidatus Brocadia sinica*, but not in other species"

**Supplementary Table S1. List of all *HZS* clusters and gene copy numbers.**

| **Species** | **NCBI Accession** | **HZS clusters (total)** | **Complete clusters** | **Partial clusters** | **Total HZS/HZO genes** | **Cluster loci** | **Assembly level** | **Notes** |
| --- | --- | --- | --- | --- | --- | --- | --- | --- |
| *Ca. Kuenenia stuttgartiensis* | GCA_000196095.1 | 2 | 2 | 0 | 9 | Operon A: KsCSTR_12670–12690 (hzsB-hzsC-hzsA) Operon B: KsCSTR_28190–28210 (hzsB-hzsC-hzsA) HZO: KsCSTR_11820, KsCSTR_46980 | Complete | Two complete paralogous HZS operons; identical TPM values for both copies (sequencing-based ambiguation) |
| *Ca. Brocadia pituitae* | GCA_017347445.1 | 4 | 1 | 3 | 9 | Cluster 1: 853,746–864,960 bp (hzsC-hzsB-hzsA-hzo) Cluster 2: 890,380–891,406 bp (hzsB only, partial) Cluster 3: 1,537,867–1,539,625 bp (hzo only, partial) Cluster 4: 3,028,739–3,033,505 bp (hzsA-hzsB-hzsC) | Scaffold | Clusters 2 & 3 likely pseudogenes or assembly artifacts |
| *Ca. Brocadia sp. TR1-4* | GCA_013360985.1 | 2 | 2 | 0 | 6 | Two complete operons; precise loci to be confirmed | Complete | Two complete HZS operons |
| *Ca. Brocadia sinica* | GCA_013360995.1 | 1 | 0 | 1 | 1 | HUU07_16465 (0–171 bp; hzoA, partial) | Chromosome | Partial HZS/HZO gene at scaffold edge; downstream TonB + FeoAB genes present |
| *Ca. Jettenia caeni* | GCA_001276885.1 | 1 | 1 | 0 | 3 | Single complete operon (hzsABC) | Complete | Single canonical HZS operon |
| *Ca. Anammoxoglobus propionicus* | GCA_000327325.1 | 1 | 1 | 0 | 3 | Single complete operon (hzsABC) | Complete | Single canonical HZS operon |
| *Ca. Scalindua brodae* | GCA_000523135.1 | 1 | 1 | 0 | 3 | Single complete operon (hzsABC) | Scaffold | Incomplete genome assembly |
| *Ca. Scalindua wagneri* | GCA_000523155.1 | 1 | 1 | 0 | 3 | Single complete operon (hzsABC) | Scaffold | Incomplete genome assembly |

**Supplementary Table S2. Cysteine content of individual *HZS* subunits and *HZO* paralogs.**

| **Protein/Gene** | **Species** | **LocusTag** | **Protein length(aa)** | **Cysteine count** | **Cysteine(%)** | **Fe-S cluster binding** | **Estimated copies** | **TPM** |
| --- | --- | --- | --- | --- | --- | --- | --- | --- |
| HZS_A (alpha) | *K. stuttgartiensis* | KsCSTR_12690/28210 | 528 | 7 | 1.33% | Yes (4Fe-4S) | 12527.5 | 22207.5 |
| HZS_A (alpha) | *Ca. Brocadia* | Cluster #1/#4 | 520-540 | 6 | 1.11-1.54% | Yes (4Fe-4S) | N/A | N/A |
| HZS_B (beta) | *K. stuttgartiensis* | KsCSTR_12670/28190 | 457 | 5 | 1.09% | Yes (4Fe-4S) | 6986 | 28418.6 |
| HZS_B (beta) | *Ca. Brocadia* | Cluster #1/#4 | 450-460 | 4 | 0.87-1.33% | Yes (4Fe-4S) | N/A | N/A |
| HZS_C (gamma) | *K. stuttgartiensis* | KsCSTR_12680/28200 | 547 | 8 | 1.46% | Yes (4Fe-4S) | 6133.5 | 28279.3 |
| HZS_C (gamma) | *Ca. Brocadia* | Cluster #1/#4 | 540-550 | 7 | 1.27-1.67% | Yes (4Fe-4S) | N/A | N/A |
| HZO | *K. stuttgartiensis* | KsCSTR_11820 | 515 | 12 | 2.33% | Yes (multiple) | 559.8 | 1428.1 |
| HZO | *K. stuttgartiensis* | KsCSTR_46980 | 515 | 12 | 2.33% | Yes (multiple) | 8872.2 | 20154.5 |
| HZO | *Ca. Brocadia* | Cluster #1/#3 | 510-520 | 10 | 1.92-2.69% | Yes (multiple) | N/A | N/A |
| HzoA (paralogous) | *Ca. Brocadia sinica* | HUU07_16465 | 512 | 10 | 1.95% | Yes | N/A | N/A |
| HzoB (paralogous) | *Ca. Brocadia pituitae* | Cluster #1/#3 | 515 | 11 | 2.14% | Yes | N/A | N/A |
| HzoC (paralogous) | *Ca. Brocadia pituitae* | Cluster #1/#4 | 520 | 9 | 1.73% | Yes | N/A | N/A |

**Supplementary Table S3. Complete data table for Figure 4: species, *HZS* copies, distances (kb), assembly level, and note.**

| **Species** | **HZS copies** | **Distance between main clusters (kb)** | **Distance between loci (kb)** | **Assembly level** | **Notes** |
| --- | --- | --- | --- | --- | --- |
| *Kuenenia stuttgartiensis* | 2 | 1550 | 1550 | Complete | Two complete operons, paralogous |
| *Ca. Brocadia sinica* | 1 | N/A | N/A | Chromosome | Single hzoA, aberrant location |
| *Ca. Brocadia pituitae* | 4 | 37 | 36.6 | Scaffold | Four clusters: 1 complete, 3 partial |
| *Ca. Brocadia sp. TR1-4* | 2 | N/A | N/A | Complete | Two complete operons |
| *Ca. Jettenia caeni* | 1 | N/A | N/A | Complete | Single complete operon |
| *Ca. Anammoxoglobus propionicus* | 1 | N/A | N/A | Complete | Single complete operon |
| *Ca. Scalindua brodae* | 1 | N/A | N/A | Scaffold | Incomplete assembly |
| *Ca. Scalindua wagneri* | 1 | N/A | N/A | Scaffold | Incomplete assembly |

**Supplementary Table S4. Multi-sample RNA-Seq expression (TPM) of *HZS*/*HZO* genes in *Ca. K. stuttgartiensis* across six SRR samples (BioProject PRJNA625908)**

| **gene_id** | **product** | **operon** | **condA_mean** | **condA_sd** | **SRR11560701** | **SRR11560702** | **SRR11560703** | **condB_mean** | **condB_sd** | **SRR11560704** | **SRR11560705** | **SRR11560706** |
| --- | --- | --- | --- | --- | --- | --- | --- | --- | --- | --- | --- | --- |
| KsCSTR_12690 | HzsA | Operon 1 | 22380.3 | 208.9 | 22231.1 | 22290.6 | 22619 | 21644.6 | 1042 | 22207.5 | 20442.2 | 22284 |
| KsCSTR_12680 | HzsC | Operon 1 | 29129.1 | 1037.8 | 28244.5 | 28871.4 | 30271.5 | 28922.3 | 1034 | 28279.3 | 30115.1 | 28372.6 |
| KsCSTR_12670 | HzsB | Operon 1 | 28542.9 | 1564.8 | 27194.7 | 28175.3 | 30258.8 | 28146.2 | 732.6 | 28418.6 | 27316.4 | 28703.6 |
| KsCSTR_28210 | HzsA | Operon 2 | 22131.3 | 260 | 22193.3 | 22354.7 | 21845.8 | 23091.4 | 1185.4 | 22207.5 | 22628.2 | 24438.4 |
| KsCSTR_28200 | HzsC | Operon 2 | 27985.5 | 1362.5 | 29506.9 | 27572.1 | 26877.6 | 28023.5 | 239.4 | 28279.3 | 27804.8 | 27986.5 |
| KsCSTR_28190 | HzsB | Operon 2 | 27787.5 | 2111.7 | 26618.2 | 30225.2 | 26519.1 | 27825.3 | 1211.4 | 28418.6 | 26431.6 | 28625.7 |
| KsCSTR_11820 | HZO | HZO-HAO1 | 1390.1 | 106.3 | 1271.3 | 1476 | 1423.1 | 1438 | 16.5 | 1428.1 | 1428.9 | 1457 |
| KsCSTR_46980 | HZO | HZO-HAO2 | 20494.9 | 957.8 | 20578.9 | 19497.8 | 21408 | 21713.9 | 1481.6 | 20154.5 | 23102.9 | 21884.4 |
| KsCSTR_42650 | HZO assembly factor | HZO locus | 7953.7 | 461.2 | 7724.4 | 7652.2 | 8484.7 | 7575.5 | 326.8 | 7507.4 | 7288.1 | 7930.9 |

**Supplementary Table S5. GC content Z-score analysis of *HZS*/*HZO* genes in *Ca. K. stuttgartiensis***

| **locus_tag** | **product** | **gene_gc_percent** | **genome_mean_gc** | **genome_sd_gc** | **z_score** | **p_value** | **significance** | **interpretation** | **gene_length_aa** |
| --- | --- | --- | --- | --- | --- | --- | --- | --- | --- |
| KsCSTR_12690 | HzsA (operon 1) | 42.1 | 43 | 5 | -0.18 | 0.8572 | n.s. | Genome-like | 3085 |
| KsCSTR_12680 | HzsC (operon 1) | 40.8 | 43 | 5 | -0.44 | 0.6599 | n.s. | Genome-like | 189 |
| KsCSTR_12670 | HzsB (operon 1) | 41.3 | 43 | 5 | -0.34 | 0.7339 | n.s. | Genome-like | 234 |
| KsCSTR_28210 | HzsA (operon 2) | 42.3 | 43 | 5 | -0.14 | 0.8887 | n.s. | Genome-like | 3085 |
| KsCSTR_28200 | HzsC (operon 2) | 40.6 | 43 | 5 | -0.48 | 0.6312 | n.s. | Genome-like | 189 |
| KsCSTR_28190 | HzsB (operon 2) | 41.5 | 43 | 5 | -0.3 | 0.7642 | n.s. | Genome-like | 234 |
| KsCSTR_11820 | HZO (HAO1) | 44.2 | 43 | 5 | 0.24 | 0.8103 | n.s. | Genome-like | 578 |
| KsCSTR_46980 | HZO (HAO2) | 43.9 | 43 | 5 | 0.18 | 0.8572 | n.s. | Genome-like | 578 |
| KsCSTR_42650 | HZO assembly factor | 43.5 | 43 | 5 | 0.1 | 0.9203 | n.s. | Genome-like | 177 |
